# Mass spectrometry analysis of newly emerging coronavirus HCoV-19 spike S protein and human ACE2 reveals camouflaging glycans and unique post-translational modifications

**DOI:** 10.1101/2020.04.29.068098

**Authors:** Zeyu Sun, Keyi Ren, Xing Zhang, Jinghua Chen, Zhengyi Jiang, Jing Jiang, Feiyang Ji, Xiaoxi Ouyang, Lanjuan Li

## Abstract

The pneumonia-causing COVID-19 pandemia has prompt worldwide efforts to understand its biological and clinical traits of newly identified HCoV-19 virus. In this study, post-translational modification (PTM) of recombinant HCoV-19 S and hACE2 were characterized by LC-MSMS. We revealed that both proteins were highly decorated with specific proportions of N-glycan subtypes. Out of 21 possible glycosites in HCoV-19 S protein, 20 were confirmed completely occupied by N-glycans, with oligomannose glycans being the most abundant type. All 7 possible glycosylation sites in hACE2 were completely occupied mainly by complex type N-glycans. However, we showed that glycosylation did not directly contribute to the binding affinity between SARS-CoV spike protein and hACE2. Additionally, we also identified multiple sites methylated in both proteins, and multiple prolines in hACE2 were converted to hydroxylproline. Refined structural models were built by adding N-glycan and PTMs to recently published cryo-EM structure of the HCoV-19 S and hACE2 generated with glycosylation sites in the vicinity of binding surface. The PTM and glycan maps of both HCoV-19 S and hACE2 provide additional structural details to study mechanisms underlying host attachment, immune response mediated by S protein and hACE2, as well as knowledge to develop remedies and vaccines desperately needed nowadays.

## Introduction

The on-going coronavirus pandemia since December 2019 (COVID-19) has now become a global health emergence (Dong et al., 2020; Wu et al., 2020b; Zhou et al., 2020b). The disease is caused by a newly identified beta-coronavirus HCoV-19 (also known as SARS-CoV-2) which is closely related to severe acute respiratory syndrome coronavirus (SARS-CoV). Like the SARS, the HCoV-19 causes lower respiratory tract infection (LRTI) which may eventually develop to atypical pneumonia that requires intensive care and mechanical ventilation (Guan et al., 2020; Wang et al., 2020; Zhou et al., 2020a). Compared to SARS-CoV, the HCoV-19 is more contagious and is infecting more population worldwide thus causing substantially more casualties (Guan et al., 2020; Zhou et al., 2020a). Although possible viral reservoirs including bats and pangolin have been suggested for HCoV-19 (Zhang et al., 2020; Zhou et al., 2020b), more investigations are need to ascertain its source and routes of zoonotic transmission (Wu et al., 2020a; Xu, 2020; Zhang and Holmes, 2020). To date, no approved vaccine or remedy specific for coronaviruses infection, including that of HCoV-19, is available.

Akin to the SARS-CoV, the virion surface spike glycoprotein encoded by HCoV-19 S gene is essential for target cell attachment and fusion processes (Hoffmann et al., 2020; Walls et al., 2020a; Wan et al., 2020). Given the importance of S protein in COVID-19 pathogenesis and its potential immunogenicity for vaccine development, global efforts have been made to elucidate the structure of S protein shortly after the publishing of the first HCoV-19 sequence (Walls et al., 2020a; Wrapp et al., 2020; Wu et al., 2020b). The HCoV-19 S glycoprotein is a 1273-amino acid precursor polypeptide and can be cleaved by host cell proteases (cathepsin L, TMPRSS2) into S1 fragment that contains the receptor binding domain (RBD) to attach host receptor human Angiotensin I Converting Enzyme 2 (hACE2), and the S2 fragment responsible for the subsequent membrane fusion (Hoffmann et al., 2020; Walls et al., 2020a; Wu et al., 2020b). The S protein is predicted to have a cleavable N-terminal signal sequence (1-15), which presumably directs the protein toward the endoplasmic reticulum (ER) for extensive glycan decoration before virion packing.

Glycosylation is one of the most prominent post-translational modifications (PTM) in many viral spike or envelop proteins, and has been shown to mediate host attachment, immune response as well as virion packaging and budding (Chang and Zaia, 2019; de Groot, 2006; Fukushi et al., 2012; Li et al., 2017; Parsons et al., 2019; Raman et al., 2016; Shih et al., 2006; York et al., 2019; Zheng et al., 2018; Zhou et al., 2010). In particular, the binding of coronavirus S proteins to their respective receptors has been shown to be mediated by its oligomannose N-glycan (Li et al., 2017; Parsons et al., 2019; Zheng et al., 2018). Additionally, C-type lectin DC-SIGN and L-SIGN can enhance viral entry via their binding to the S protein glycans (Han et al., 2007; Jeffers et al., 2004). Therefore, numerous antiviral strategies have been designed to interfere protein glycosylation or glycan-based interaction (Shih et al., 2006; Vincent et al., 2005; Zheng et al., 2020; Zhou et al., 2010), and glycosylation is considered as a key aspect to develop effective vaccine (Chen et al., 2014; Kumar et al., 2020).

Like the SARS-CoV spike protein which contains 23 N-linked glycosylation sequons (N-X-S/T, X≠P), HCoV-19 spike protein is predicted to host 22 per protomer or 66 per trimer (Walls et al., 2020a). However, the potential glycosites pattern in S1 of HCoV-19 S protein is different to that of SARS-CoV, while the glycosylation sites in S2 region are significant conserved between HCoV-19 and SARS-CoV. The difference of S1 glycosylation pattern may link to biological and clinical characteristics of HCoV-19 which is profoundly different to other coronavirus. In this study, we report a comprehensive N-glycosylation profile, as well as other PTMs, of HCoV-19 S protein and hACE2 elucidated by high resolution mass spectrometry analyses, based on which reinterpretation of current HCoV-19 S protein structural model was made to highlight important glycan features related to COVID-19 pathogenesis. Nonetheless, we demonstrate that the binding of HCoV-19 S protein and hACE2 is independent on their glycosylation status.

## Methods

### Expression and purification of HCoV-19 spike ectodomain and hACE2

HCoV-19 S gene (virus isolate: Wuhan Hu-1; GenBank number QHD43416.1) was synthesized (Genscript) with codons optimized for insect cell expression. Its ectodomain (Val16-Pro1213) was cloned into pFastBac vector (Life Technologies Inc.) with a N-terminal honeybee melittin signal peptide and C-terminal His6 and Flag tags. HCoV-19 S protein was expressed in Sf9 insect cells using the Bac-to-Bac system (Life Technologies Inc.) and harvested from cell culture medium followed by purification procedure using Ni-NTA column and Superdex 200 gel filtration column (GE Healthcare) in tandem. The extracellular domain of hACE2 (Gln18-Ser740, NP_068576.1) with C-terminal Fc tag was expressed in HEK 293 cells were purified by protein A sepharose beads (GE Healthcare).

### SDS-PAGE analysis

To test their glycosylation status of HCoV-19 spike and hACE2 protein, both proteins were deglycosylated by PNGase F (NEB, 1:50) overnight at 37°C in PBS. Both the glycosylated and deglycosylated forms of HCoV-19 spike and hACE2 protein were then analyzed by 15% polyacrylamide gel electrophoresis (SDS-PAGE) followed by Coomassie blue staining.

### Glycopeptide sample preparation

Proteins were first digested into tryptic peptides according to (Sun et al., 2016). Briefly, hACE2 and S protein were first deduced by 10 mM dithiothreitol in 50 mM ammonium bicarbonate (ABC) for 45 min at room temperature and then alkylated by iodoacetamide for 45 mins at room temperature in the dark. Proteins were then cleaned up by acetone precipitation and resuspended in 50 mM ABC. The alkylated glycoproteins were then digested for 14 h at 37°C using sequencing-grade trypsin, chymotrypsin or endoproteinase Lys-C, all purchased from Promega, with a protein:protease ratio of 1:50 in 50 mM ABC. The peptides were desalted using HLB columns (Waters).

The peptide samples were separated into duplicates. The first duplicate was deglycosylated by PNGase F (NEB, 1:100) overnight at 37°C in 50 mM ABC prepared in pure H_2_O^18^. The second duplicate was used to enrich intact N-glycopeptides by hydrophilic interaction liquid chromatography (HILIC). Briefly, the GlycoWorks (Waters) cartridges were pre-conditioned with loading buffer comprised of 15 mM ammonium acetate (AmA) and 0.1% trifluoroacetic acid (TFA) in 80% Acetonitrile (ACN). Peptides in loading buffer were applied to the cartridges and unbound flow-through (FT) fraction was collected. The column was washed two times with loading buffer. Glycopeptides were eluted in 0.1% TFA. All peptide fractions were desalted by house-made C18 stagetips before LC-MSMS analyses.

### LC-MSMS experiment

About 500 ng of peptides were analysis using an Ultimate 3000 nanoflow liquid chromatography system (Thermo Scientific, USA) connected to a hybrid Q-Exactive HFX mass spectrometry (Thermo Scientific, USA). The mass spectrometer was operated in data-dependent mode with a full scan MS spectra followed by MS2 scans recording the top 20 most intense precursors sequentially isolated for fragmentation using high energy collision dissociation (HCD). The MS and MS/MS spectra were recorded using the Xcalibur software 2.3 (Thermo Scientific, USA). Detail parameters for LC separation and mass spectrometry acquisition can be found in Supplementary Table 1 and 2, respectively.

### Bioinformatics

The acquired MS raw files from deglycosylated peptides were searched by MaxQuant (version 1.6.10.43) against the human ACE2 sequence from UniProtKB and HCoV-19 S sequence (YP_009724390.1_3) from NCBI. Cysteine carbamidomethylation was set as fixed modification, while methionine oxidation and O^18^ deamidation on glutamic acid were set as variable modification. Trypsin with up to two missed cleavages was set. Mass tolerance of 15 ppm and 4.5 ppm were set for first and main search, respectively. For comprehensive PTM analysis, phosphorylation (S/T/Y), acetylation (K), methylation (K/R/E), succinylation (K), crotonylation (K), farnesylation (K/Nterm), myristoylation (K/Nterm), palmitoylation or prenylation, glycosylphosphatidylinositol and oxidation on proline were individually investigated in parallel database searches for respective variable modification. Peptide level 1% FDR was set to filter the result. Confident identification of PTM was based on localization probability of 99%. Site occupancy for each PTM was calculated by dividing the peak intensities of the modified peptides (MP) and corresponding non-modified peptides (NP) using the equation: MP/(MP+NP)*100.

To investigate N-glycosylation forms, acquired MS raw files from HILIC experiments were searched by pGlyco (version 2.2.2) (Liu et al., 2017) against the same hACE2 and HCoV-19 S sequences. Cysteine carbamidomethylation was set as fixed modification, while methionine oxidation was set as variable modification. Trypsin with up to two missed cleavages was set. Mass tolerance of 5 ppm and 15 ppm were set for precursor and fragment mass tolerance, respectively. Potential glycan fragments within MS2 spectra were annotated by built-in pGlyco. gdb glycan structure database (Liu et al., 2017). Precursor intensity of each glycopeptide was extracted by MaxQuant feature detection algorithm.

### Structural model refinement

Glycan structure in pdb format was downloaded or predicted by GLYCAM-Web (https://dev.glycam.org/gp). The N-linked glycan models of hACE2 and trimeric spike protein were made by manually adding the MS-identified glycan at each site based on previous models (PDB codes 6M18 for hACE2 and 6VXX for spike protein) within Coot (Emsley et al., 2010). The modified residues within each model were generated with Coot (Emsley et al., 2010). In addition, model of HCoV-19 S protein and hACE2 complex (PDB 6M0J) were used to shown location of PTM surrounding the binding area. All structural figures were performed by PyMOL (Schrödinger Inc. U.S.A.).

### Evaluation of binding of HCoV-19 S protein and hACE2 by bio-layer interferometry (BLI)

Binding of HCoV-19 S protein and hACE2 was measured on an Octet Re96E (ForteBio) interferometry system. Briefly, HCoV-19 S protein were immobilized using aminopropylsilan biosensors (18-5045, ForteBio). To evaluate the influence of glycosylation of HCoV-19 S on binding, S protein were first incubated with either 1000U/mL PNGase F (NEB) or deactivated PNGase F at 37 °C for 14 h. After washing by kinetic buffer (1x PBS, pH 7.4, 0.01% BSA, and 0.002% Tween 20), the association between HCoV-19 S and hACE2 was measured for 180 seconds at 30°C by exposing sensors to hACE2 in kinetic buffer at concentration of 12.5, 25, 50, 100, 200 nM, respectively. After binding phase, the sensors were exposed to 1x kinetic buffer for dissociation for 300 seconds at 30°C. Signal baseline was subtracted before data fitting using the equimolar binding model. Mean k_on_, k_off_ values were determined with a global fit model using all data. A parallel experiment was also performed using hACE2 loaded sensor incubated with HCoV-19 S protein solution. Analogously, immobilized hACE2 was pretreated by either PNGase F or deactivated PNGase F to evaluate the influence of hACE2 glycosylation on binding. In all experiment, PNGase F was deactivated by heating at 75 °C for 10 min.

## Results

### Determination of N-linked glycosylation sites on HCoV-19 S protein and hACE2

As shown in (Figure 1A), PNGase F deglycosylation resulted in deceasing molecular weight of both HCoV-19 S protein and hACE2 on SDS-PAGE. Digestion by PNGase F releases N-linked glycans from Asn within the N-X-S/T motif, which also resulted in Asn deamidation by losing one hydrogen and one nitrogen while incorporating one oxygen derived from the solvent (Bailey et al., 2012). To perform a thorough survey of glycosylated sites, protease digested peptides from both proteins were subjected to PNGase F deglycosylation in H_2_O^18^ that enables incorporation of O^18^ and leads to a +2.98 Da mass increment, which was used to mark glycosylated sites.

**Figure 1.**
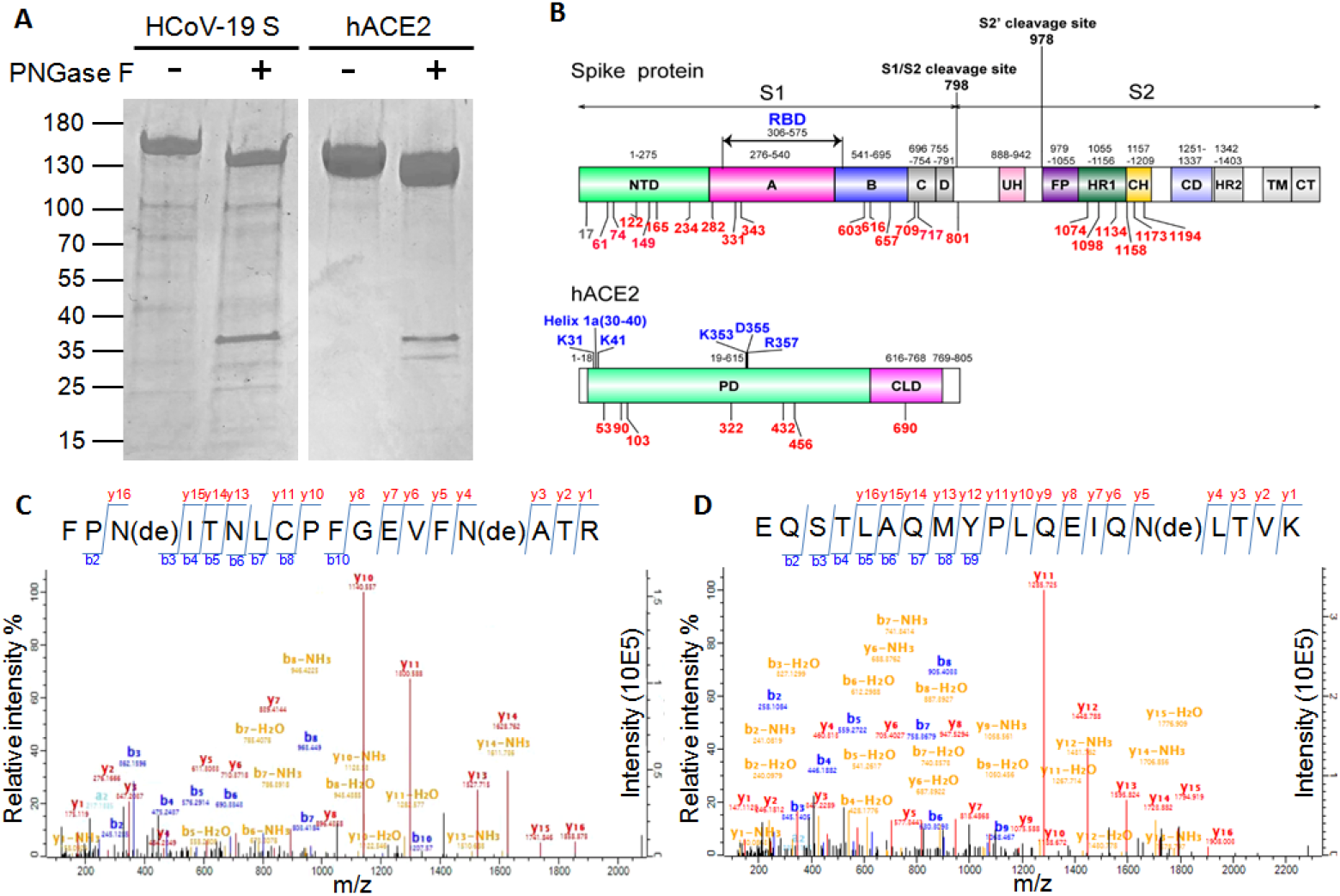
Potential glycosylation sites in HCoV-19 spike protein and hACE2. **A**: 15% SDS-PAGE analysis of intact and deglycosylated form of HCoV-19 spike protein and hACE2. Molecular weight markers are shown on the left. **B**: schematic representation of functional subunits and domains of HCoV-19 spike protein (upper panel) and hACE2 (lower panel). CD, connector domain; CH, central helix; CT, cytoplasmic tail; FP, fusion peptide; TM, transmembrane domain; UH, upstream helix; HR1/2, heptad repeat 1/2. Blue indicated sites possibly responsible for interaction between the S protein and hACE2. Potential glycosylation sites within in each domain were listed in braces. Red indicated identified glycosylated sites in this study. **M**ass spectra of identified deglycopeptide containing N331 and N343 in HCoV-19 spike protein (**C**) and deglycopeptide containing N90 in hACE2 (**D**).

Analysis of the deglycosylated peptides from S protein by LC-MS/MS confirmed 20 N-linked glycosylation sites, including N61, N74, N122, N165, N234, N282, N331, N343, N603, N616, N657, N709, N717, N801, N1074, N1098, N1134, N1158, N1173 and N1194 (Table 1). Two remaining N-X-S/T sites, the N17 and N149 were not identified in any deglycosylated peptide. However, peptide with glycosylated N149 was identified directly without PNGase F treatment as described later, therefore leading to a total of 21 glycosylation sites out of 22 potential sites with N-X-S/T sequons in HCoV-19 S protein (Figure 1B). In addition, quantitative analysis of site occupancy showed there were 18 sites completely glycosylated, while N603, N657 reached 43% and 74% occupancy (Table 1). Example mass spectra showing evidence of N331 and N343 glycosylation was shown in (Figure 1C).

**Table 1.**
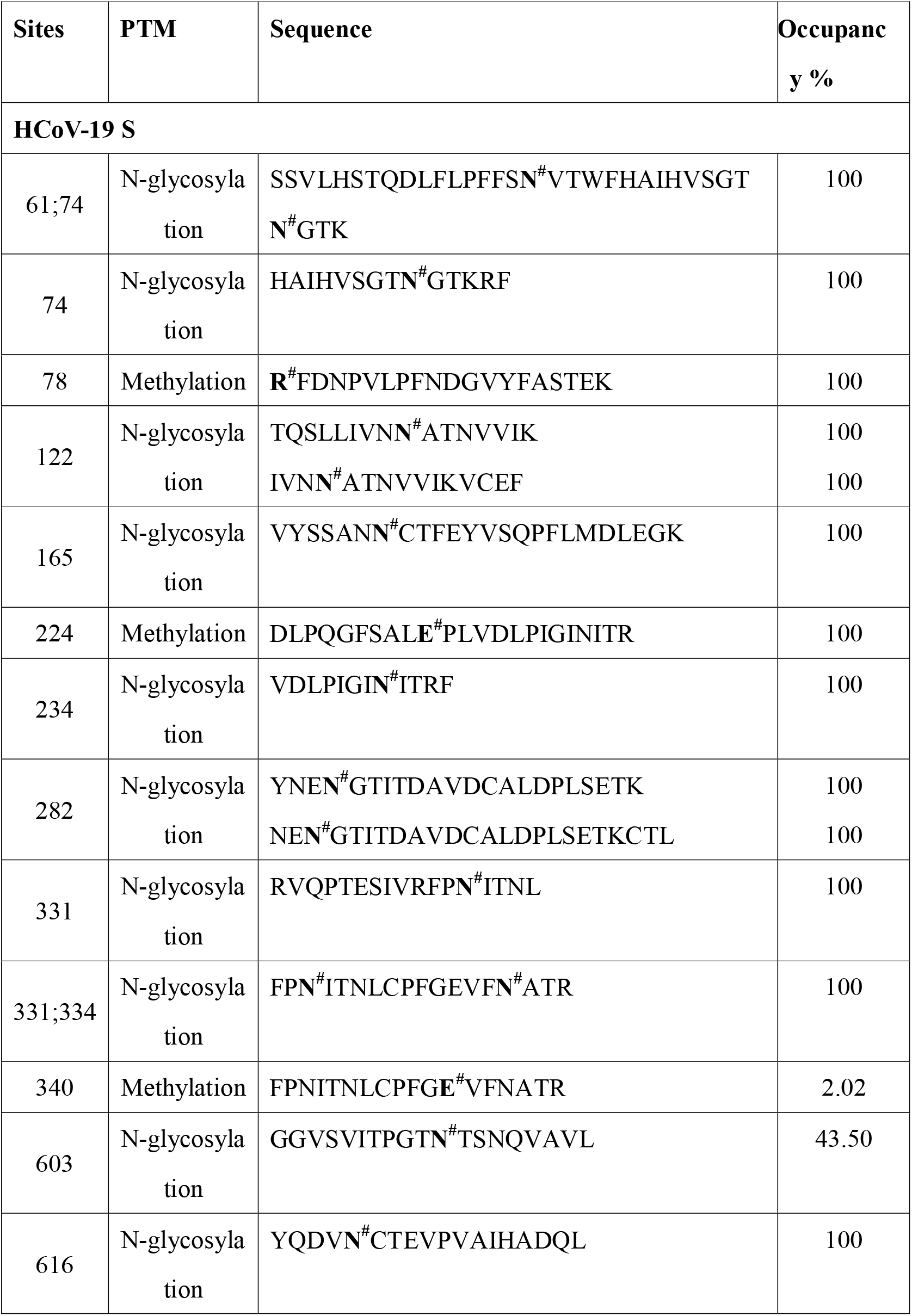
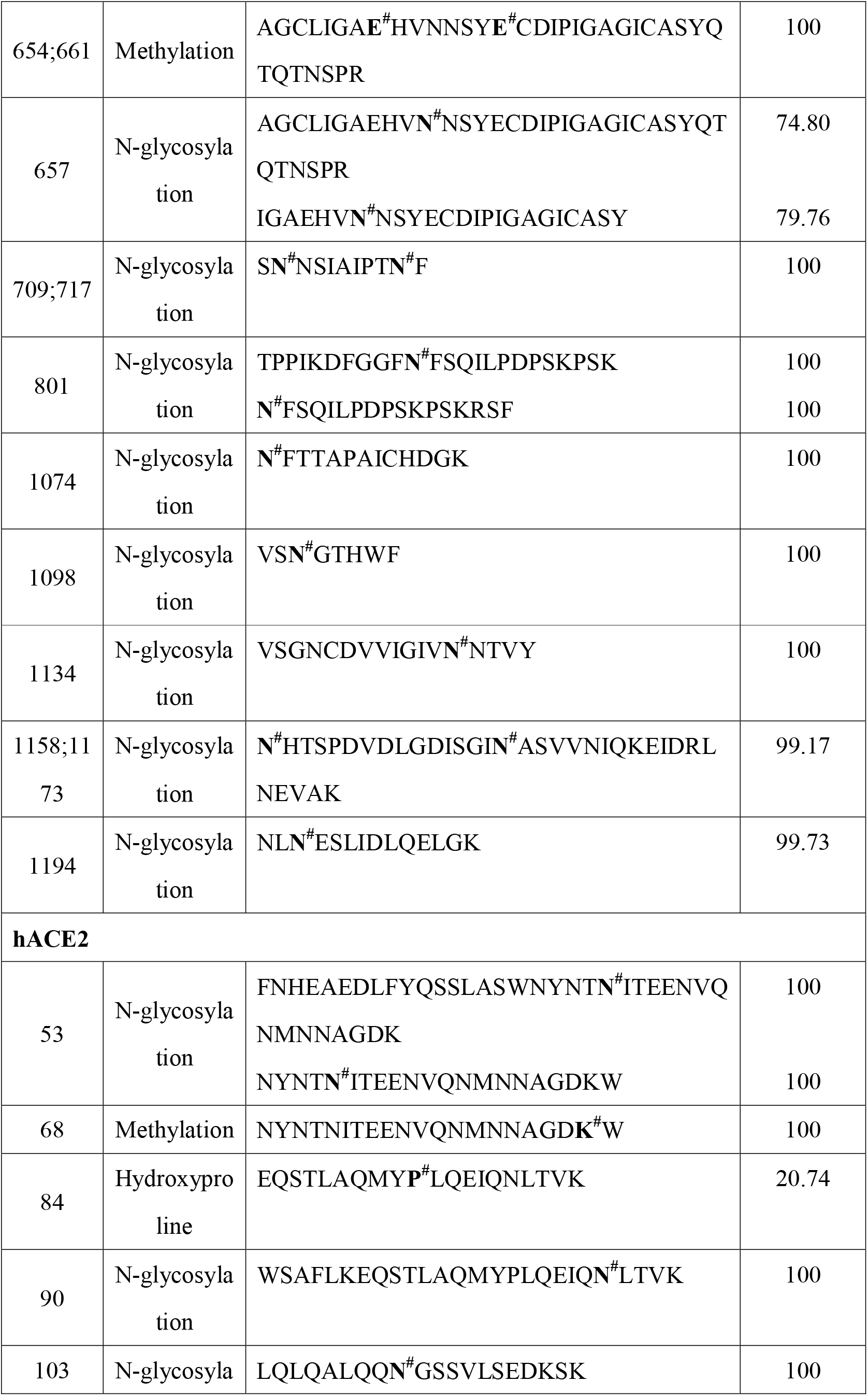
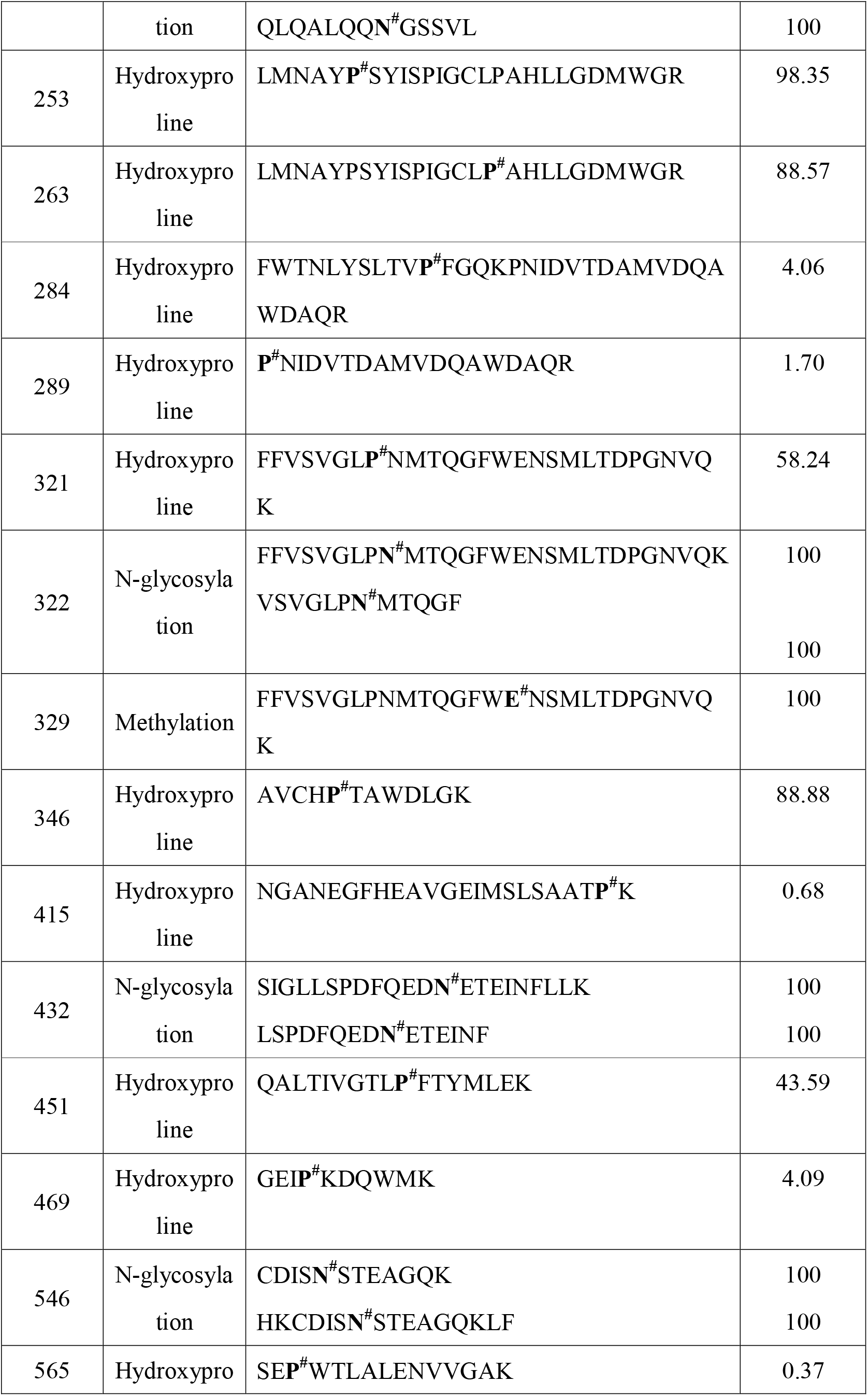
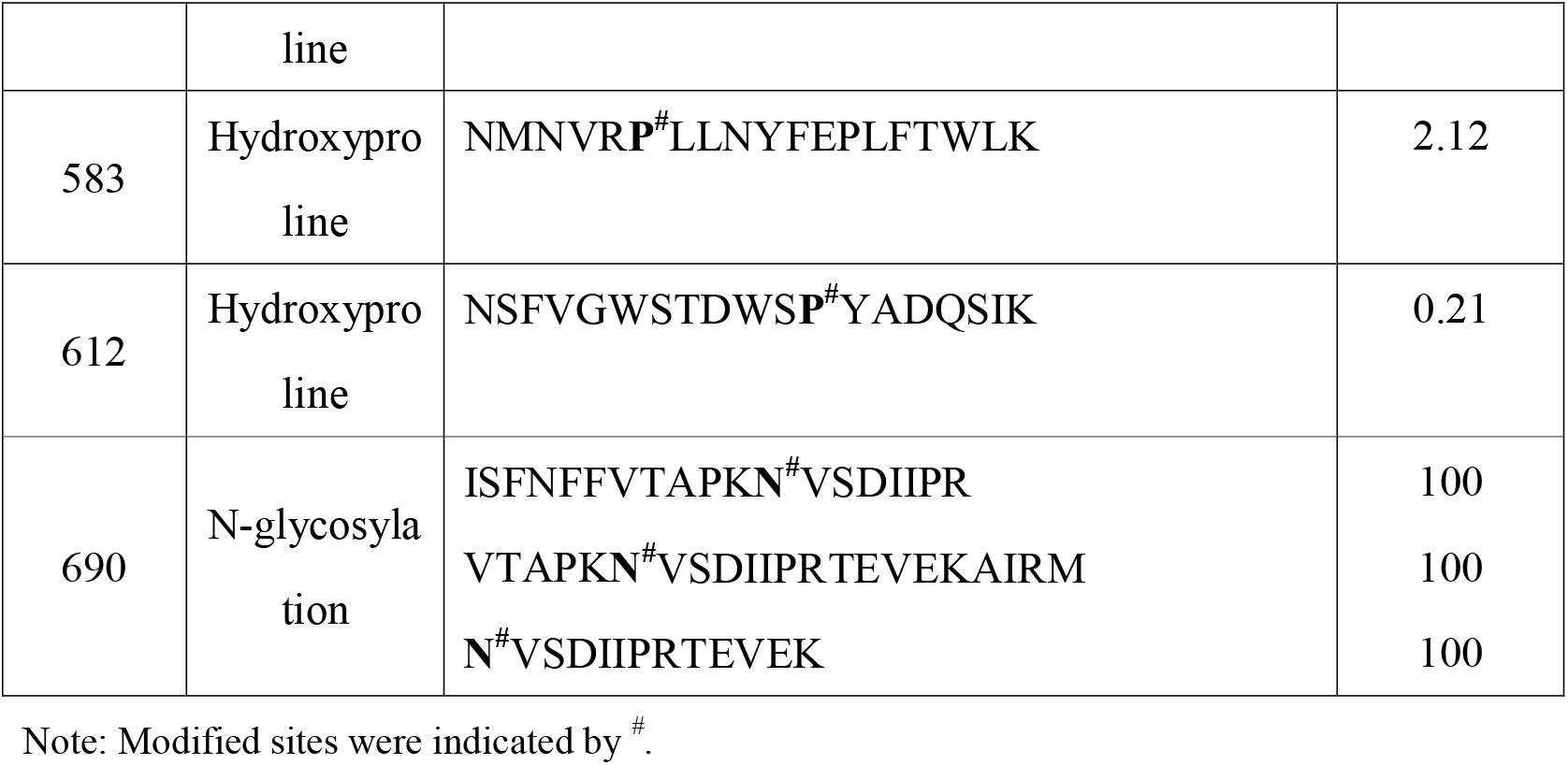
Summary of PTM identified in deglycosylated peptides from HCoV-19 S and hACE2 protein.

All 7 possible glycosylation sites in hACE2, N53, N90, N103, N322, N432, N546 and N690 were confirmed as glycosylated in our experiments (Table 1). As example, mass spectra of N90 glycosylation were shown in Fig 1D. Quantitative analysis of site occupancy showed all 7 sites were completely glycosylated (Table 1). All these data suggested that both HCoV-19 S protein and hACE2 are highly decorated by N-glycans.

### Identification of PTMs of HCoV-19 S protein and hACE2

To unveil possible post-translational modification (PTM) other than glycosylation, LC-MSMS data of deglycosylated peptides from both HCoV-19 S protein and hACE2 were subjected to several rounds of spectra-database matching for common PTMs. The major PTM found in both proteins was methylation on lysine, arginine or glutamic acid, as summarized in Table 1. Using occupancy >50% as criteria to identify site dominantly decorated with PTMs, we found 78R, 224E, 654E and 661E in HCoV-19 S protein, 57E, 68K and 329E in hACE2 is highly methylated. In addition, proline at site 253, 263, 321 and 346 in hACE2 can be found prominently oxidized and converted to hydroxyproline. However, common PTMs such as phosphorylation, acetylation or other acylations were not found in HCoV-19 S protein and hACE2.

### Global N-glycosylation profile of HCoV-19 S protein and hACE2

To resolve glycan camouflage on surface of HCoV-19 S protein and hACE2, intact glycopeptides derived from protease digestion and fractionated by HILIC SPE were directly subject to LC-MSMS analysis specifically designed to detect peptides with extra molecular weight due to N-glycan attachment. Due to the highly heterogeneous nature of glycan chain, in our study a unique glycopeptides was designated as combination of both unique peptide sequence with a specific N-glycan composition. According to this criteria, 419 and 467 unique N-glycopeptides were identified from HCoV-19 S proteins and hACE2, respectively (Table 2 and 3). Out of 20 HCoV-19 S glycosylated sides identified by PNGase F experiment, 19 were confirmed by intact glycopeptide profiling. Despite all 7 glycosylation sites in hACE2 were identified previously in deglycosylated peptides labeled with deamidation, N-glycan profile was obtained in 5 sites: N90, N103, N432, N546 and N690. Example N-glycopeptides spectra from HCoV-19 S and hACE2 could be found in Supplementary Figure 1.

**Table 2.**
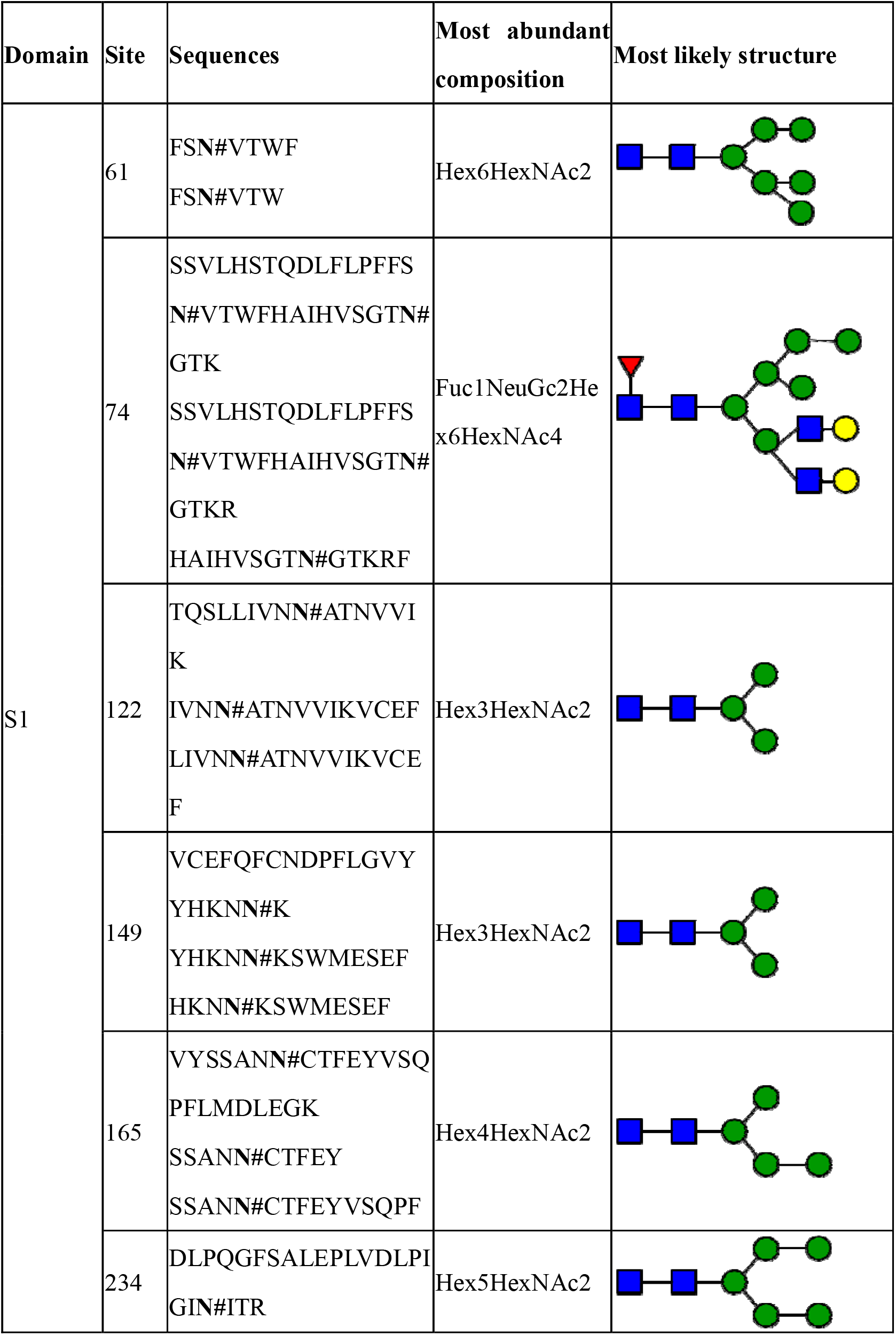
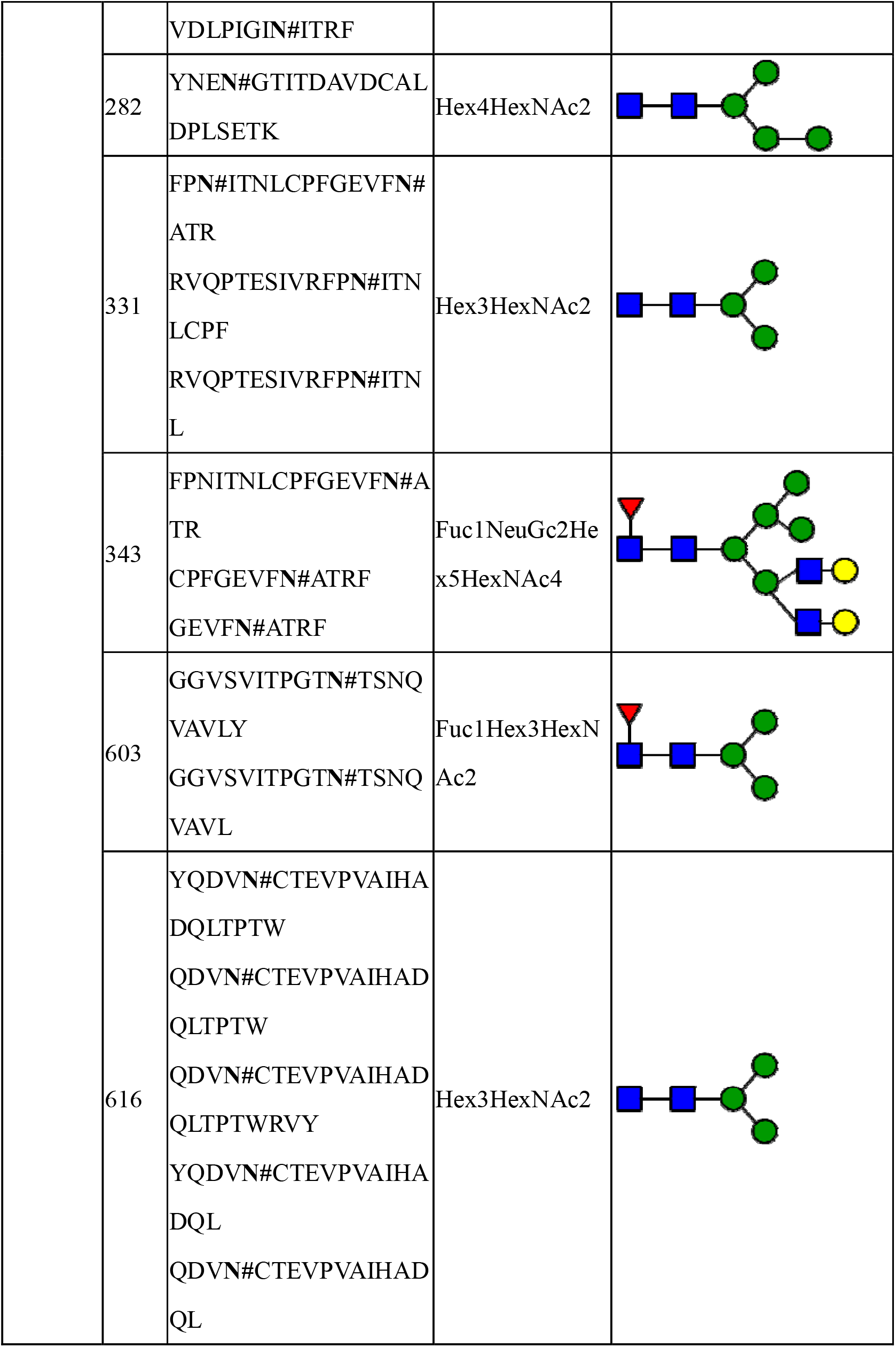
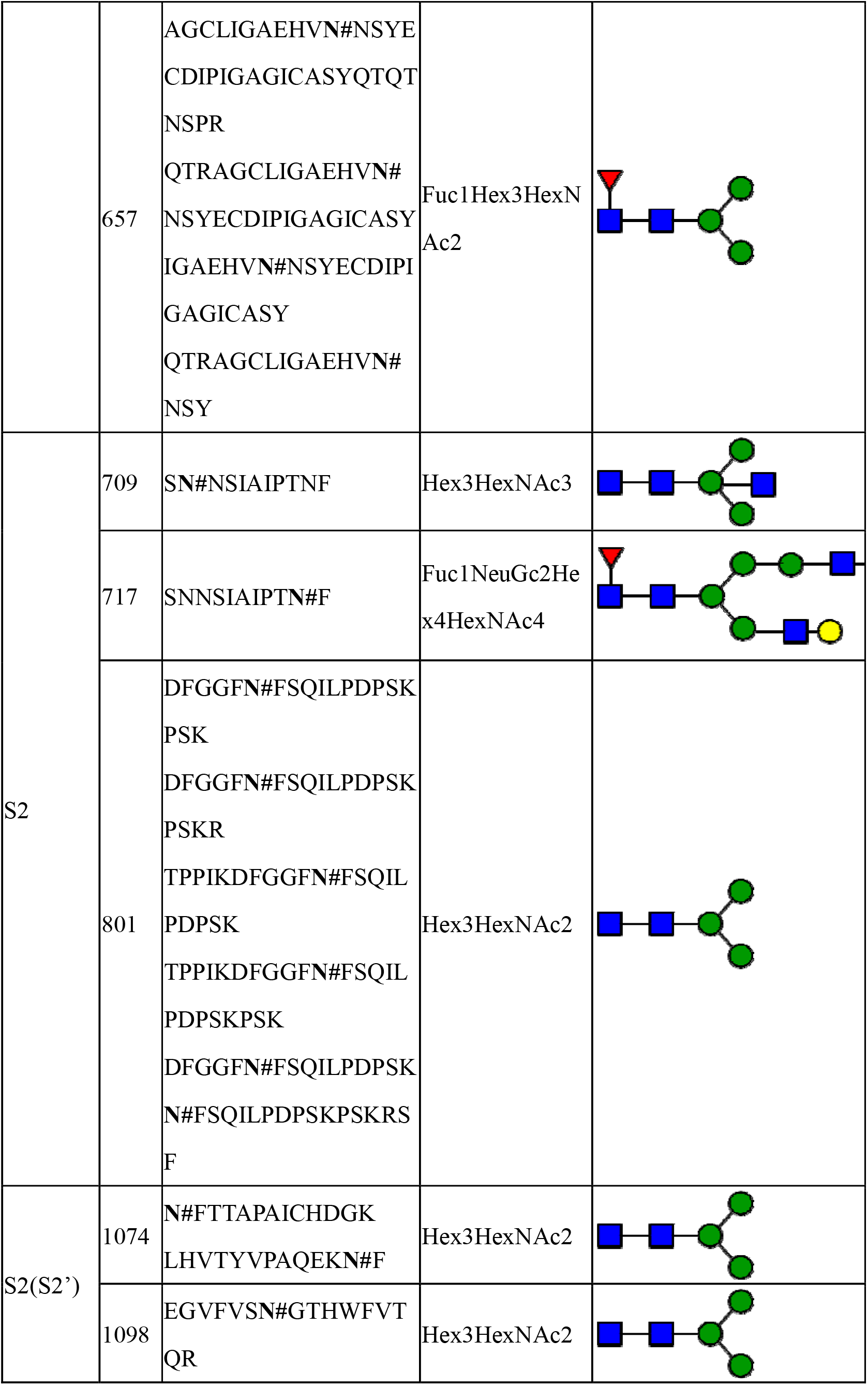
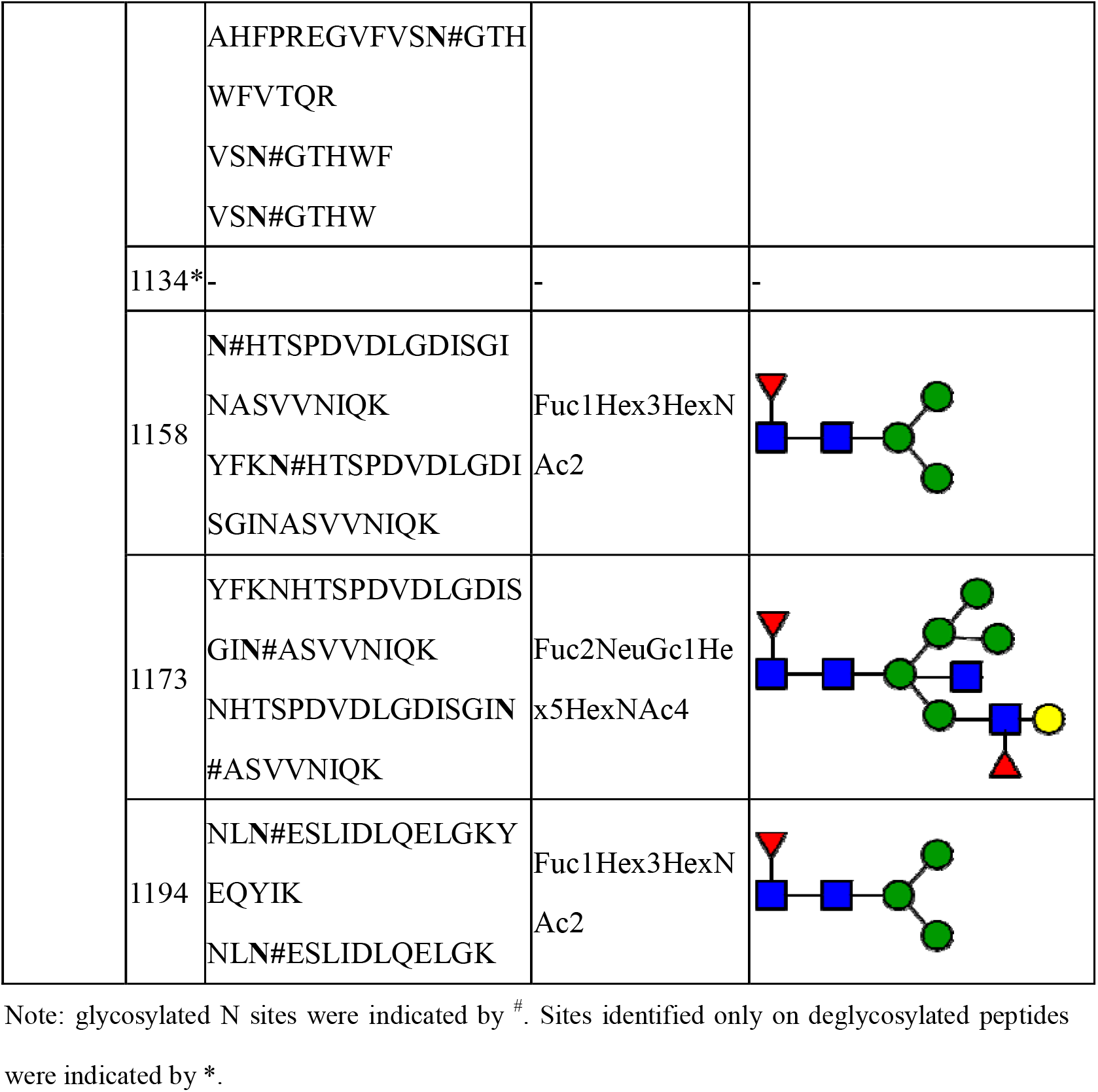
Summary of glycopeptide and N-glycan identification in HCoV-19 S protein.

**Table 3.**
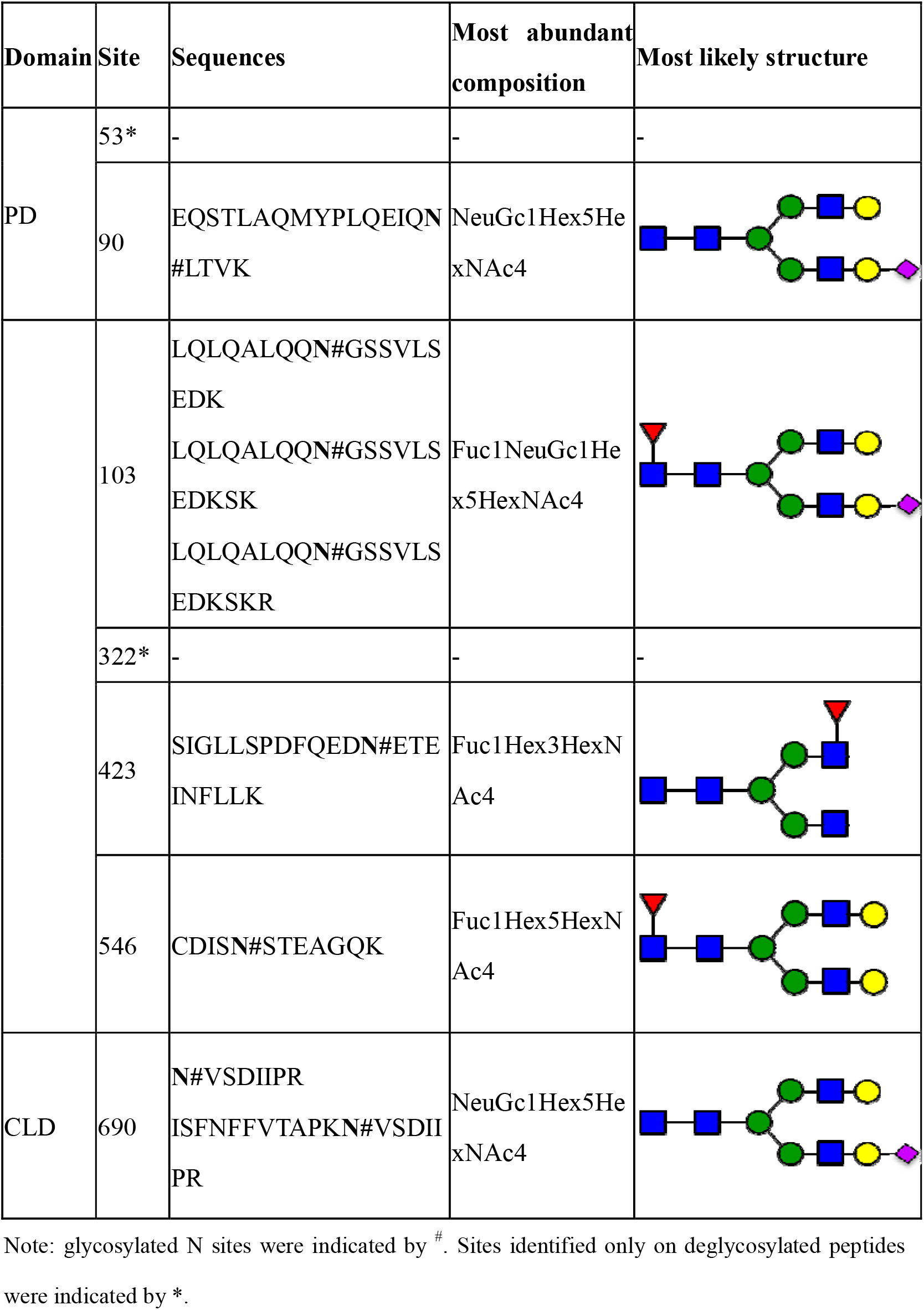
Summary of glycopeptide and N-glycan identification in hACE2 protein.

A total of 144 N-glycans were found in HCoV-19 S protein, with majority of them containing the common N-acetylglucosamine core (Table 2). All N-glycosites in S protein were attached with multiple types of N-glycans, with N343 decorated by the most diverse N-glycans. The HCoV-19 S N-glycan composition preferentially comprising pauci- or high-mannose type oligosaccharides, except in four N-glycosites (N73, N343, N717 and N1173) containing a high proportion of complex and hybrid N-glycans (Figure 3A, 3B and Table 2). By LC-MS intensity, pauci-mannose Hex3HexNAc2 was the most common N-glycan in HCoV-19 S protein. Interestingly, there was no evidence of sialic acid component in the N-glycan of HCoV-19 S protein.

**Figure. 2.**
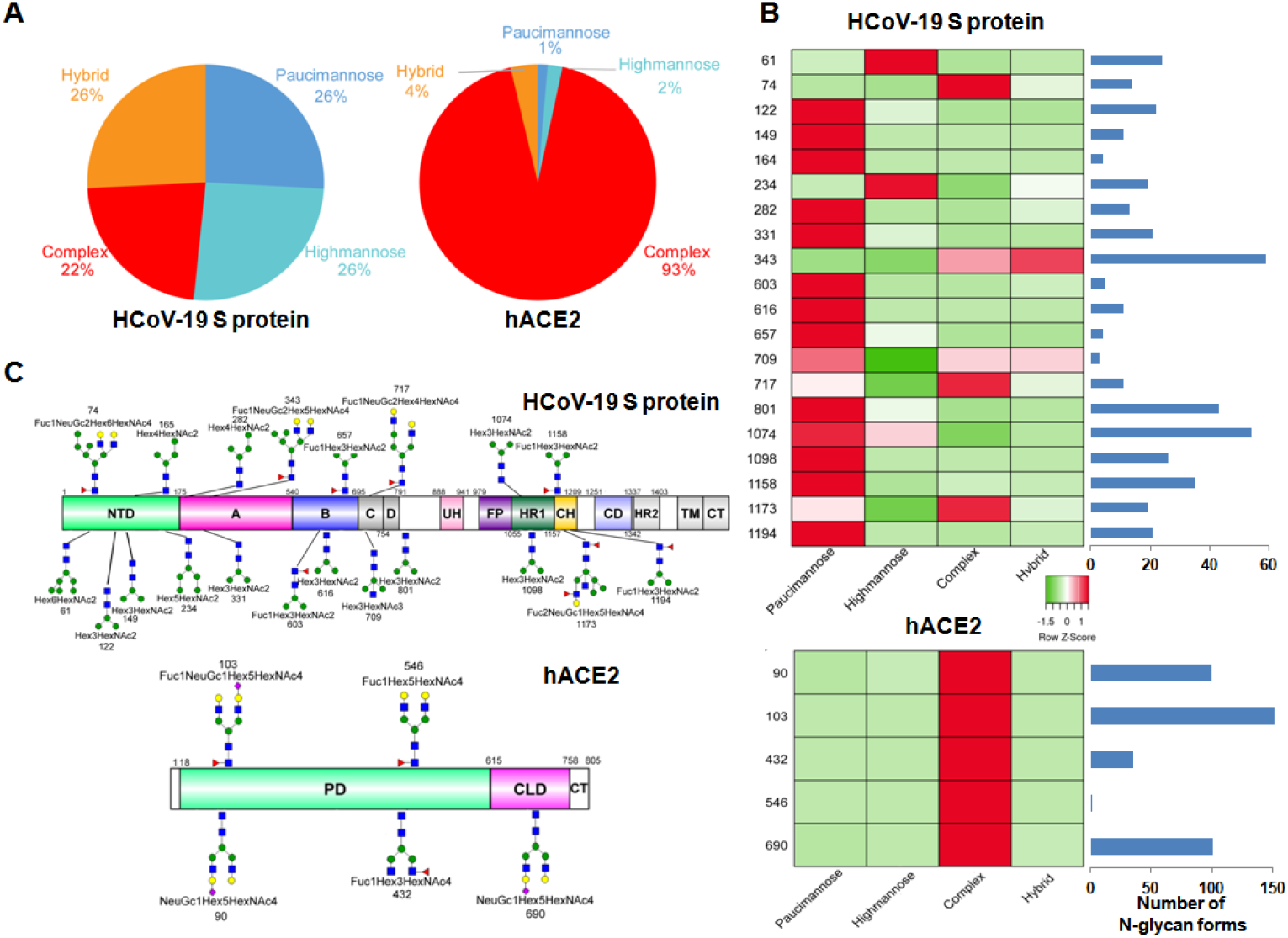
Summary of N-glycopeptide survey in HCoV-19 Spike protein and hACE2. **A**: Overall percentages of 4 major N-glycan categories identified in either protein. **B:** schematic representation of predominant N-glycan forms in each site related to functional domains in either protein. **C:** relative LC-MS intensity of 4 major N-glycan categories identified in each site were presented in heatmap, alongside with total number of identified glycans summarized in bars plots.

**Figure. 3.**
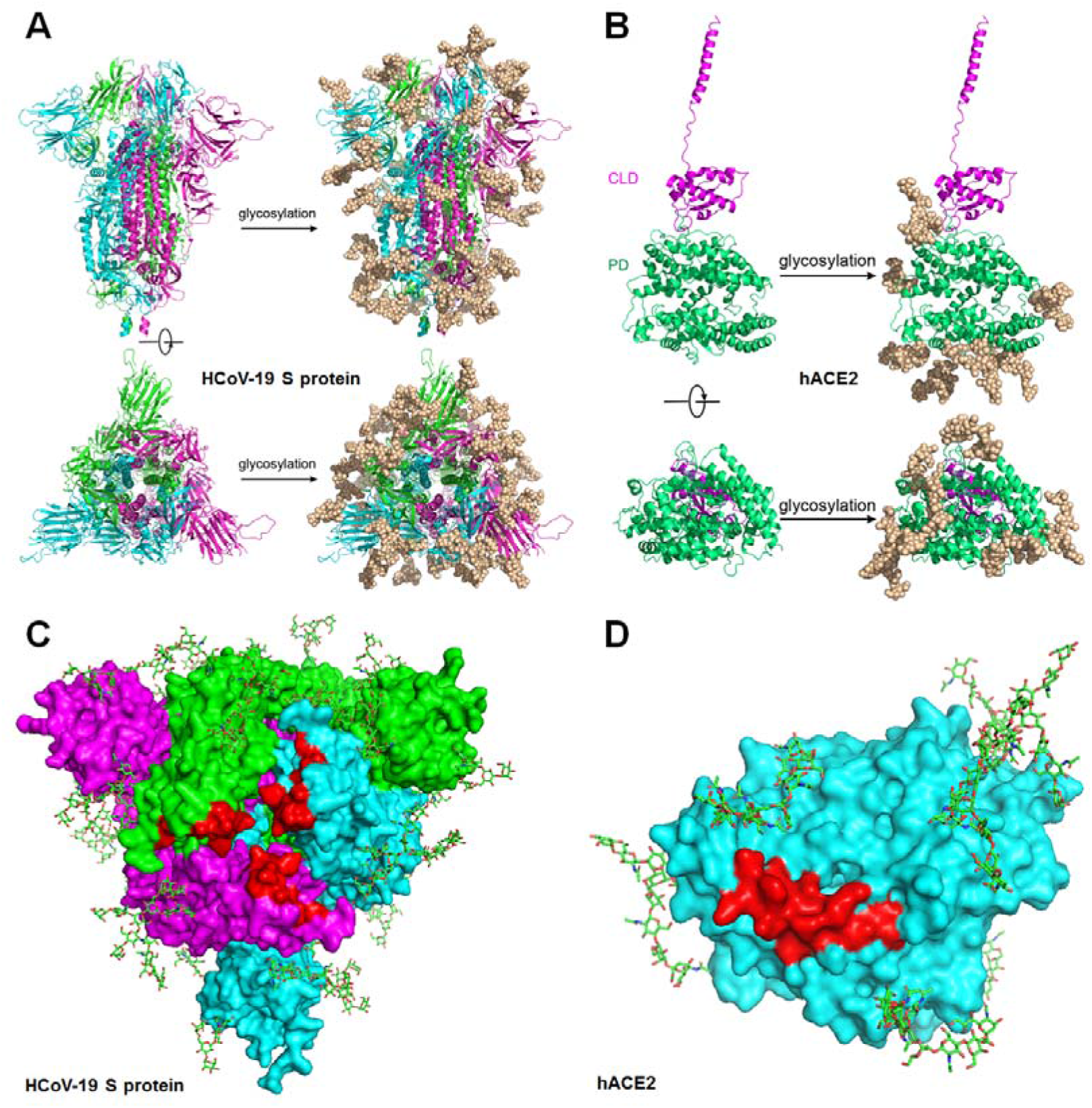
Refined structure model of HCoV-19 Spike trimer and hACE2 incorporating N-glycans. 3D ribbon diagrams of HCoV-19 spike trimer colored by protomer (A) and hACE2 colored by major domains (B). For both model, side-view (upper panel) and top-view looking towards the viral or cellular membrane (lower panel) are shown, model without (left panel) or with glycans (right panel) were both provided. Diagrams showing binding sites (red) with glycans in vicinity on the surface of HCoV-19 spike trimer (C) and hACE2 (D) in top-view. Glycans were presented in sticks.

A total of 220 N-glycans was found in hACE2, with 78.2% of them being complex type (Figure 3A), while hybrid, high-mannose and pauci-mannose glycan only constitute 10.9%, 5.0% and 5.9%, respectively (Figure 3B and Table 3). By LC-MS intensity, the most dominant hACE2 glycan was NeuGc1Hex5HexNAc4 which was predicted to be a bi-antennary complex containing a sialic acid end. N-glycan profile in all hACE2 sites was primarily complex type. It is notable that N90, N103 and N690 all contain more than 100 types of N-glycans, with their most dominant N-glycan form all containing sialic acid.

Collectively, our LC-MSMS data confirms that both HCoV-19 spike protein and its receptor hACE2 are indeed heavily N-glycosylated at most of its predicted N-X-S/T sequon. Summary of most dominant N-glycan composition and predicted structure in HCoV-19 spike protein and hACE2 were illustrated in Figure 3C.

### Structure modeling of HCoV-19 S-hACE2 complex refined with glycan and PTM details

By adding chemical structures of the most abundant N-glycans in each site based on LC-MSMS results to the most updated cyro-EM model of HCoV-19 s protein and hACE2, we generated atomic models that represented the most likely spatial distribution of the N-glycans on both proteins (Fig. 3A and 3B). Despite 4 sites (N74, N149, N1158, and N1194) of spike protein were not shown in the model for the missing residues in model 6VXX, our model suggested the camouflaging N-glycans shielded more than 2/3 of the HCoV-19 S protein surface, which could potentially lead to host attachment and immune evasion. Additionally, both HCoV-19 s protein and hACE2 model showed the glycans at N331, N343 of HCoV-19 s protein and N90 of hACE2 were in the proximity of, albeit not exactly inside of, the binding area of both proteins (Fig. 3C and 3D). Model of S-hACE2 complex also showed that 3 methylation sites in hACE2 (57E, 68K and 329E) formed a trident structure which enclosed the contact area formed between K353-R357 of hACE2 and N501 of HCoV-19 S (Supplementary Figure 2).

### Binding of HCoV-19 S protein and hACE2 does not depends on N-glycosylation

To understand the contribution of protein glycosylation to the interaction HCoV-19 S protein with hACE2, we compared the binding kinetics and affinity of the purified hACE2 ectodomain to glycosylated and deglycosylated HCoV-19 S-ECD immobilized at the surface of biosensors in a BLI experiment. We found that hACE2 bound to HCoV-19 S pretreated with active or inactive PNGase F with comparable equilibrium dissociation constants Kd of 1.7 nM (Figure 4A) and 1.5 nM (Figure 4B), respectively. The affinity determined in this study for hACE2 binding to HCoV-19 S is in line with (Walls et al., 2020a). Comparable affinity was also observed when HCoV-19 S ECD was bound to the immobilized hACE2 (Kd = 16.7 nM, Figure 4C) or deglycosylated hACE2 (Kd = 18.2 nM, Figure 4D). Detailed summary of BLI binding kinetics can be found in Supplementary Table 3.

**Figure. 4.**
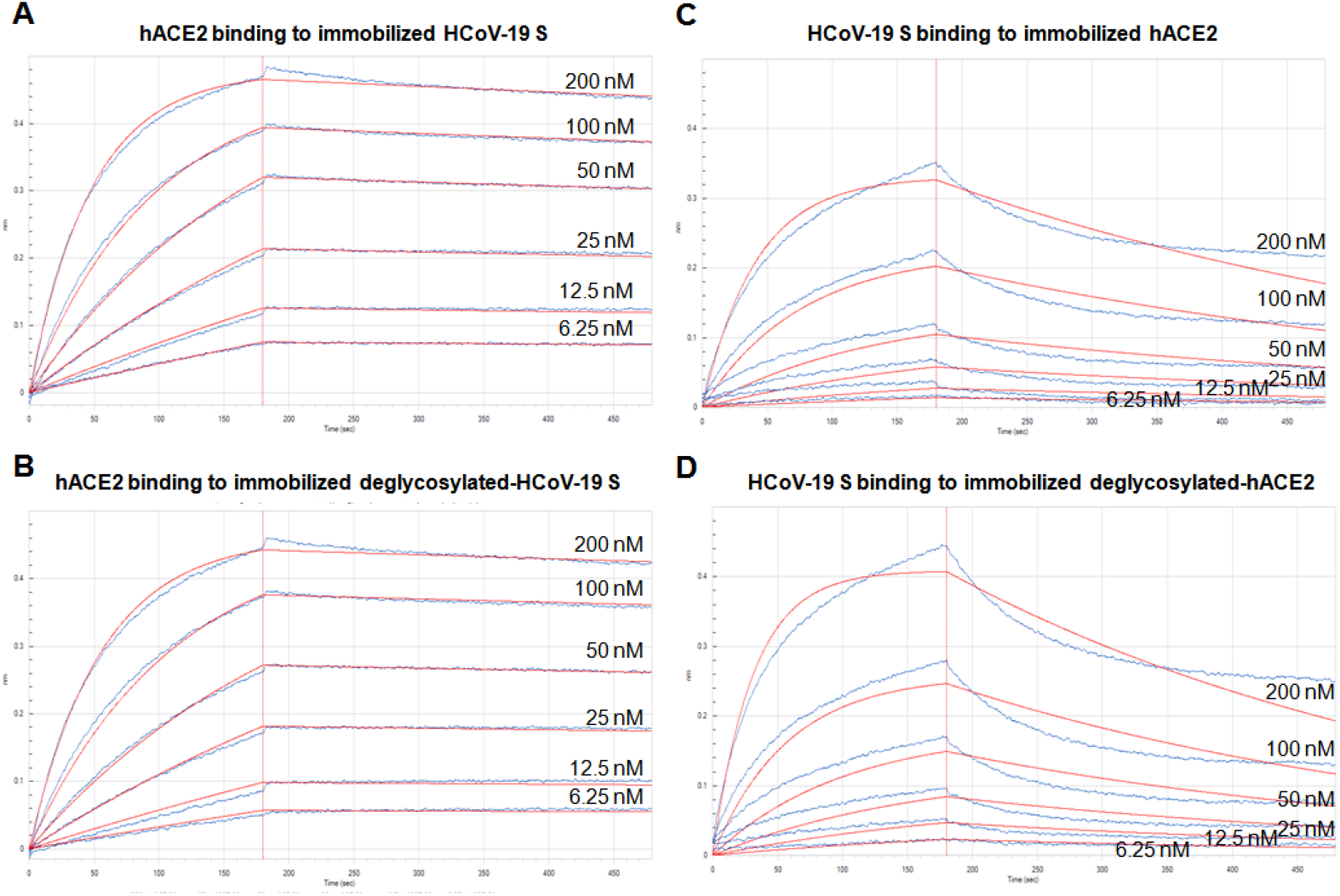
Impact of glycosylation on binding between HCoV-19 Spike protein and hACE2. Binding of HCoV-19 S protein and hACE2 was measured by biolayer interferometry. Biosensors with immobilized intact HCoV-19 S protein (A) and its deglycosylated form (B) were exposed to hACE2 at concentration of 12.5, 25, 50, 100, 200 nM. Swap experiments were conducted using biosensors with immobilized intact hACE2 (C) and its deglycosylated form (D) exposed to HCoV-19 S protein at concentration of 12.5, 25, 50, 100, 200 nM. Red lines correspond to a global fit of the data using an equimolar binding model.

## Discussion

Glycosylation is a ubiquitous and complex PTM that greatly extend the structural and functional diversity for many proteins. However, comprehensive characterization of protein N-glycosylation is technically challenging. In this study, deglycosylated peptides which are easier to be analyzed by LC-MS than their glycosylated precursors, were profiled to confirmed N-glycosylated sites in HCoV-19 S and hACE2. The O^18^ deamidation modification significantly boost the confidence for glycosite identifications as the isotope was incorporated by PNGase F mediated hydrolysis in H_2_O^18^. Intact N-glycopeptide analyses were then performed to further verify the glycosites, and provided detailed information about the compositional and structural heterogeneity of N-glycans associated with each site. Our investigation showed that all sites except N17 were highly glycosylated in HCoV-19 S protein. According to the N-linked glycan model of spike protein, a large proportion of the protein surface is covered by glycans, which is consistent with that in previous reports (Walls et al., 2020b; Wrapp et al., 2020). When compared spike proteins from HCoV-19 and SARS-CoV, it is noticeable that the majority of differences of glycosylation sites occurs in the distal S1 subunit, therefore leading to significant difference in glycan profile in the outermost canopy of the virus formed by spike trimer clusters. Alteration of glycosites might be the results of host or environmental selective pressure, and could implicate dramatic difference in viral infectivity, pathogenesis and host responses.

The N glycans in S protein are markedly heterogeneous, which greatly extends conformational flexibility and epitope diversity. While a small proportion of complex and hybrid N-glycans were found in S protein, most sites were occupied by oligomannose glycans, which is in line with previous report on SARS-CoV (Krokhin et al., 2003; Song et al., 2004; Ying et al., 2004). Many viral proteins are hallmarked by high level of oligomannose-type glycans, probably as a sign of incomplete glycan maturation due to high glycosite density that results in steric hindrance (Watanabe et al., 2019). Interestingly, N343, the glycosite closet to the binding surface, is decorated by the most diverse N-glycans and primarily by hybrid and complex forms. Moreover, like in case of SARS-CoV and Marburg viral proteins (Feldmann et al., 1994; Ritchie et al., 2010), we found that sialic acid incorporation in SARS-CoV spike protein glycans is negligible. Regarding the ongoing global efforts to generate vaccine, which primarily targets S protein as the candidate antigen, the diverse glycan forms decorating large section of S trimer surface should be considered by vaccine developers as they could drastically modulate the protein immunogenicity. We should also expect that vaccines generated based on S proteoforms with different glycosylation status could render varying efficacy against future infection.

The binding of coronavirus S proteins to their respective receptors has been shown to be mediated by its oligomannose N-glycan (Li et al., 2017; Parsons et al., 2019; Zheng et al., 2018). SARS-CoV spike protein contains 3 RBD-associated glycosites (N318, N330 and N357), while HCoV-19 only has 2 (N331 and N343) that surrounds the binding pocket. In contrary to our postulation that N-glycan at these two sites might contribute polar interaction to receptor binding, the BLI binding assay suggested deglycosylation did not change the affinity of SARS-CoV spike protein to hACE2. However, this negative BLI binding results do not exclude the possibility that glycosylation could affect other viral entry steps, including protease cleavage, and glycans mediated interactions with DC/L-SIGN which were documented to enhance viral entry in SARS-CoV (Han et al., 2007; Jeffers et al., 2004). Further investigations are required to ascertain the role of glycosylation regarding the infectivity of HCoV-19.

Antigen glycosylation greatly determines host immune responses. One of the prominent consequence of viral envelope or surface proteins is immune evasion by shield off immunogenic epitopes (Raman et al., 2016; Vigerust and Shepherd, 2007; Yang et al., 2020). The complete occupancy of glycans in most glycosite of HCoV-19 S protein suggest the virus is able to invade the host in a stealth fashion. Successful innate immune evasion by the HCoV-19 at the early stage of infection might explain the long asymptomatic incubation period during which transmissible virion are produced (Guan et al., 2020; Wang et al., 2020). Cases of HCoV-19 reinfection were also reported suggesting the virus is capable to escape from antibody-mediated neutralization (Biswas et al., 2020). On the other hand, protein glycosylation can also facilitate immune response by sensor molecules or antibodies specific for glycan recognition. N-glycans are known ligands for galectins, which triggers inflammation events such as cytokine release, immune cell infiltration (Robinson et al., 2019; Wang et al., 2019), of which may contribute to the HCoV-19 pathogenesis and correlated with the disease severity. Moreover, epidemiological studies have shown that ABO polymorphism are linked to different susceptibility to both HCoV-19 and SARS-CoV infections (Cheng et al., 2005; Zhao et al., 2020). Since the ABH carbohydrate epitopes and viral protein glycans are likely to be synthesized by the same ER-resident glycosylation enzymes, anti-A or-B antibodies could partially block the viral-host interactions (Guillon et al., 2008), thereby with blood type O individuals more resistant to virus infection. It remains to be explored if the glycosylation status of S protein has implications in the inter-individual variation in terms of viral response and clinical outcome within the infected population.

The glycosylation status of hACE2 was also profiled in this study. All 7 glycosites in hACE2 were completely occupied by glycan, including the N90 in the vicinity of binding surface. However, we found the glycan did not directly contribute to the its binding with HCoV-19 S protein. This is in agreement with previous founding that the disruption of ACE2 glycosylation did not affect its binding of SARS-CoV spike protein (Zhao et al., 2015). But deglycosylated ACE2 did compromise the cellular entry and the subsequent production of infectious SARS-CoV virion (Zhao et al., 2015). Therefore, ACE2 glycosylation was still considered as an important target for intervention of coronavirus infection. In fact, the ability of chloroquine to counter HCoV-19 infection was thought to be the results of its inhibition ACE2 glycosylation in addition to its ability to increase endosomal pH level (Liu et al., 2020; Vincent et al., 2005).

Additional PTM forms other than glycosylation were also investigated in this study. Methylation was identified in several sites in both S protein and hACE2. In particular, we found 57E, 68K and 329E in hACE2 that surrounding its binding site with RBD of S protein are completed methylated. Methylation causes loss of charge and increases hydrophobicity in these sites. Four hydroxyproline (253, 263, 321, 346) were identified in the protease domain of hACE2. The extra hydroxyl group is like to increase hydrophilicity of proline in this extracellular region of hACE2. Further study is needed to investigate possible biological role of these PTMs. We did not found phosphorylation and acetylation in our dataset as previous PTM investigation on SARS-CoV also fail to identify these PTMs (Ying et al., 2004). Fatty acid acylations were also been considered in our study since they are commonly found in surface viral protein to facilitate membrane fusion and virus entry. However, our study found no evidence of acylation PTM presence in HCoV-19 S protein. Given that our multi-protease proteomic experiment provides almost full coverage for both proteins, we believe the PTMome of HCoV-19 S protein and hACE2 mainly consist of glycosylation, methylation and proline oxidation.

## Acknowledgments

We thank the proteomics and metabolomics platform in the State Key Laboratory for Diagnosis and Treatment of Infectious Diseases at Zhejiang University for glycoproteomic analysis. We thank Shuangtian Shengwu Biotech. Co.Ltd for assistance of BIL analysis. We thank pGlyco team at Institute of Computing Technology, Chinese Academy of Sciences, Beijing, China, for technical support for pGlyco analysis. We thank Dr. Ma Ping for consultation on protein expression and purification.

## Authors’ contributions

Study concept and design: ZS, LL; Samples preparation: ZS, KR, FJ, XO; LC-MSMS experiments: ZS, KR; Analysis and interpretation of data: ZS, JJ, KR; Structure Modeling: XZ, JC, ZS; Drafting of the manuscript: ZS, ZJ; Critical revision of the manuscript: LL; All authors approved the final version of the manuscript, including the authorship list.

