## Supplementary materials for "Mass spectrometry analysis of newly emerging coronavirus HCoV-19 spike S protein and human ACE2 reveals camouflaging glycans and unique post-translational modifications"

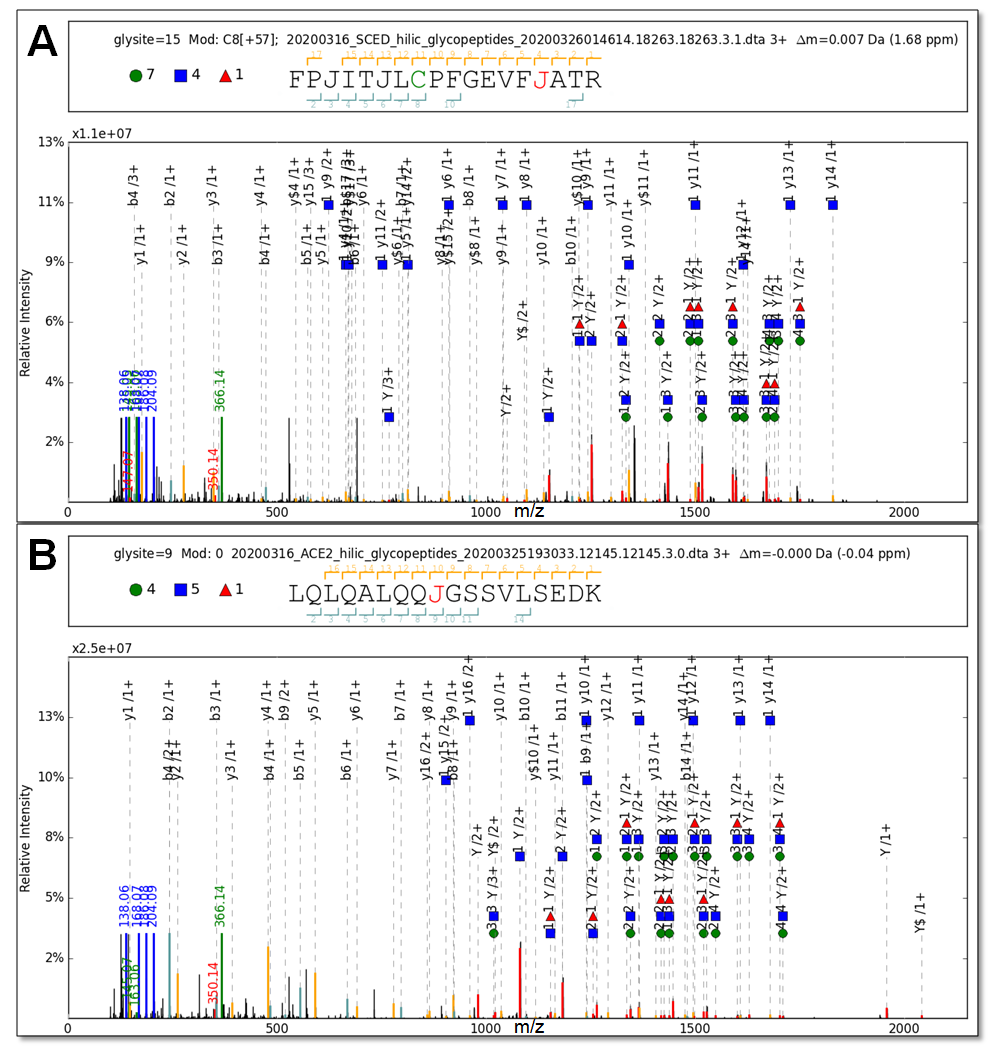

**Supplementary Figure 1**

Example mass spectra showing the identification of intact glycopeptide containing 343N glycan in S protein (A) and 103N glycan in hACE2 (B).

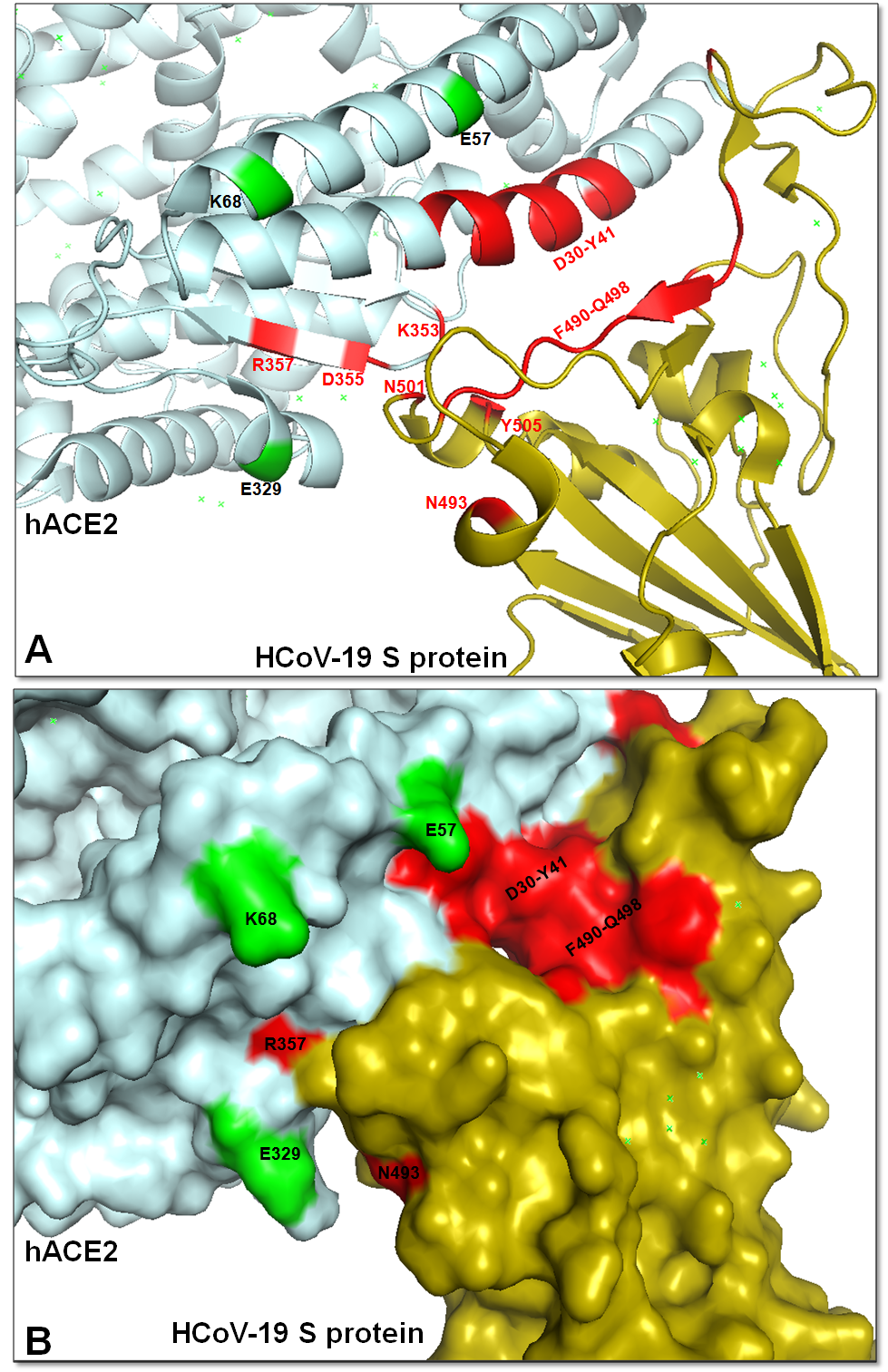

**Supplementary Figure 2**

Binding model of HCoV-19 S-protein (olive) in complexed with hACE2 (cyan) showing the backbond amino acids (A) and complex surface (B). The relevant residues in the binding site were presented in red. Methylated residues were presented in green.

**Supplementary Table 1: chromatography parameters for NanoLC-MSMS analysis**

| Column dimension | Loading:  Acclaim PepMap 100 reversed-phase pre-column (164535, Thermo Scientific), 20 mm X 75 μm, 5 μm |
| --- | --- |
|  | Analyzing:  Acclaim PepMap 100 reversed-phase column (164941, Thermo Scientific), 250 mm X 75 μm, 2 μm |
| Mobile Phase A | 2% ACN, 0.1% FA |
| Mobile Phase B | 98% ACN, 0.1% FA |
| Flow rate | 400 nL/min |
| Total LC time | 120 min |
| Gradient | 0-6 min: 3% B;  6-7 min: 3–5% B;  7-70 min: 5–18% B;  70-90 min: 18–32% B;  90-100 min: 32-80% B;  100-110 min: 80% B;  110-120 min: 80-3% B. |

**Supplementary Table 2: mass spectrometry parameters for LC-MS analysis**

|  | Peptide analysis | Glycopeptide analysis |
| --- | --- | --- |
| Electrospray voltage | 2.2 kV | |
| Total run time | 120 min | |
| MS1 resolution | 120000 | |
| MS1 AGC | 1E6 | 3E6 |
| MS1 MIT | 60 ms | 120 ms |
| MS1 range | 350-1800 Th | 800-2300 Th |
| MS2 resolution | 15000 | |
| MS2 AGC | 1E5 | 5E5 |
| MS2 MIT | 60 ms | 250 ms |
| TopN | 20 | |
| Isolation window | 1 Th | 2 Th |
| NCE | 27 | 21,28,38 |
| Min AGC target | 1E3 | |
| Charge exclusion | 1, >7, unknown | |
| Peptide match | Preferred | |
| Dynamic Exclusion | 30 s | |

**Supplementary Table 3: summary of BLI affinity assay.**

|  | **ACE2 binds to S** | **ACE2 binds to deglyco- S** | **S binds to ACE2** | **S bind to deglyco-ACE2** |
| --- | --- | --- | --- | --- |
| **Conc. (nM)** | **Response equilibrium (nm)** | | | |
| **200** | 0.4764 | 0.4602 | 0.3306 | 0.4086 |
| **100** | 0.4624 | 0.4663 | 0.2237 | 0.2579 |
| **50** | 0.5147 | 0.4809 | 0.1408 | 0.179 |
| **25** | 0.5412 | 0.5208 | 0.1003 | 0.1251 |
| **12.5** | 0.5352 | 0.4861 | 0.0599 | 0.0866 |
| **6.25** | 0.5378 | 0.485 | 0.036 | 0.0505 |
| **Kinetics summary** | | | | |
| **KD (M)** | 1.77E-09 | 1.50E-09 | 1.82E-08 | 1.67E-08 |
| **kon(1/Ms)** | 1.04E+05 | 9.00E+04 | 1.12E+05 | 1.49E+05 |
| **kdis(1/s)** | 1.85E-04 | 1.35E-04 | 2.03E-03 | 2.49E-03 |
| **Full R^2** | 0.999 | 0.9986 | 0.9826 | 0.9772 |
